# The experience of vivid autobiographical reminiscence is supported by personal semantic representations in the precuneus

**DOI:** 10.1101/197665

**Authors:** Vishnu Sreekumar, Dylan M. Nielson, Troy A. Smith, Simon J. Dennis, Per B. Sederberg

**Affiliations:** Department of Psychology, The Ohio State University, Columbus, OH, USA; Department of Psychological Science, University of North Georgia, Oakwood, GA 30566; School of Psychology, University of Melbourne, Melbourne, VIC, Australia

## Abstract

Recent studies have suggested that the human posteromedial cortex (PMC), which includes core regions of the default mode network (DMN), plays an important role in episodic memory. Whereas various roles relating to self-relevant processing and memory retrieval have been attributed to different subsystems within this broad network, the nature of representations and the functional roles they support in these brain regions remain unspecified. Here, we describe the whole-brain networks that represent *subjective*, self-relevant aspects of real-world events during autobiographical recollection. Nine participants wore a device to record images from their lives for a period of two to four weeks (lifelogging phase) and indicated the personally-salient attributes (i.e., personal semantics) of each episode by choosing multiple content tags. Two to four weeks after the lifelogging phase, participants relived their experiences in an fMRI scanner cued by images chosen from their own lives. Representational Similarity Analysis revealed a broad network, including parts of the DMN, that represented personal semantics during autobiographical reminiscence. Furthermore, within this network, the right precuneus represented personally relevant content during vivid recollection but not during non-vivid recollection. The precuneus is a hub within the DMN and has been implicated in metacognitive ability for memory retrieval. Our results suggest a more specific mechanism underlying the phenomenology of vivid reminiscence, supported by personal semantic representations in the precuneus.

## Introduction

Tulving(1993;2002) suggested that episodic memory is a unique human capability that enables us to engage in mental-time travel along a subjective timeline to reinstate past experiences. In a previous study, we identified the neural correlates of the *objective* spatiotemporal axes along which mental travel occurs during autobiographical memory retrieval (Nielson et al., 2015). However, the concept of episodic memory is incomplete without a notion of the self, the accompanying subjective dimensions of experience, and a special inwardly turned state of consciousness — termed autonoetic awareness — that guides retrieval and monitoring of autobiographical memories. In this paper, we describe the networks involved in the retrieval of the *subjective*, self-relevant content of real-world memories experienced over several weeks.

Autobiographical memory (AM) concerns our personal histories and encompasses both episodic and personal semantic memory (Conway, 2001; Levine et al., 2002). For example, knowledge about “I play ultimate frisbee every Wednesday”is part of AM, but it need not necessarily be accompanied by a specific episodic memory or vivid recollection of the details surrounding a particular instance of having played ultimate frisbee. This type of personal semantics, operationalized as autobiographical knowledge or information extracted from repeated autobiographical events, has recently garnered a lot of attention and is thought to be an intermediate entity between semantic and episodic memory (see Renoult et al., 2012, for a review). The recollective experience results only when details of a specific event are reinstated (Conway, 2001; also see Box 5 in Renoult et al., 2012).Therefore, everyday acts of memory involve guidance by retrieval of personal semantic knowledge culminating in the retrieval of a specific episode (Barsalou, 1988; Binder et al., 2009; Conway and Pleydell-Pearce, 2000). Additionally, vivid reminiscence is a hallmark of episodic recollection (Eldridge et al., 2000; Moscovitch et al., 2005) and therefore, in this study, we investigate the brain networks that subserve personal semantics and identify the specific parts of these personal semantic networks that support the phenomenological experience of vivid AM.

Given the special status of the self in AM, it is likely to engage brain networks that have previously been found to be involved in processing information in relation to the self (for a recent review, see Qin and Northoff, 2011). Specifically, the default mode network (DMN) (Buckner et al., 2008; Raichle et al., 2001) has been associated with internally oriented processing across domains like memory (Cabeza and Nyberg, 2000; Hassabis et al., 2007; Kim, 2010; Sestieri et al., 2011; Svoboda et al., 2006), prospection (Addis et al., 2007; Spreng and Grady, 2010; Spreng et al., 2009), mental imagery (Cabeza and Nyberg, 2000), and mind-wandering (Christoff et al., 2009). Consistent with this general conception of the DMN, an emerging body of neuroimaging work suggests that the human posteromedial cortex (PMC), which includes core regions of the DMN such as the retrosplenial cortex (RSc), posterior cingulate cortex (PCC) and the precuneus (pC), is involved in episodic memory (Cabeza et al., 2008; Miller et al., 2008; Rugg et al., 2002; Shannon and Buckner, 2004; Uncapher and Wagner, 2009; Vannini et al., 2011; Wagner and Davachi, 2001). Recently, attempts have been made to characterize the various subsystems of the DMN. For example, Kim (2012) proposed a dual subsystems view of the DMN where midline cortical regions including the medial prefrontal cortex (mPFC) and PCC/pC were hypothesized to be associated with self-referential processing (Hebscher et al., 2017), whereas the more lateral regions such as the MTL were thought to be involved in episodic retrieval. However, it is not clear what is represented or processed in the PMC during “self-referential processing”. It is also not known if retrieving memories of real-world experiences spanning several weeks using highly personalised visual memory cues utilizes the same networks previously identified using generic memory cues (e.g. see Vilberg and Rugg, 2008 for an argument that the observation of a left-lateralized parietal retrieval network could be a result of the limited range of verbal memory cues used in previous studies). Recently, Rissman et al. (2016) employed wearable cameras to investigate distributed brain activity patterns during memory retrieval but they focused on classifying mnemonic output (remember vs familiar vs new) rather than representational content. Therefore, critical questions remain about the specific functional roles and information content of the various regions of the parietal recollection network (Rugg and Vilberg, 2013), particularly in a relatively more ecologically valid autobiographical reminiscence task. Critically, we had access to participant-generated content labels for each recorded episode from their lives which allowed us to track specific representations of personal semantics across each individual’s brain as they relived their experiences cued by images chosen from their own lives.

In a previous study focused on the medial temporal lobe (MTL), we found that the anterior hippocampus represents objective space and time content, i.e., the “where”and “when”during retrieval of AM extending over spatiotemporal scales of up to 30 Km and 1 month (Nielson et al., 2015). In the current paper, we perform multi-variate pattern analysis on activity across the whole brain to investigate the brain networks that subserve personal semantics (i.e., the “what”of AM) and identify the specific parts of these personal semantic networks that support the phenomenological experience of vivid AM recollection.

## Results

Participants wore a lifelogging device (Figure 1A) that captured images and other sensor data as they went about their everyday activities for a period of two to four weeks. At the end of each day, participants tagged each episode with personally salient attributes chosen from a drop-down menu using a web interface (see Figure 1B, and Materials and Methods, Table 2).

**Figure 1.**
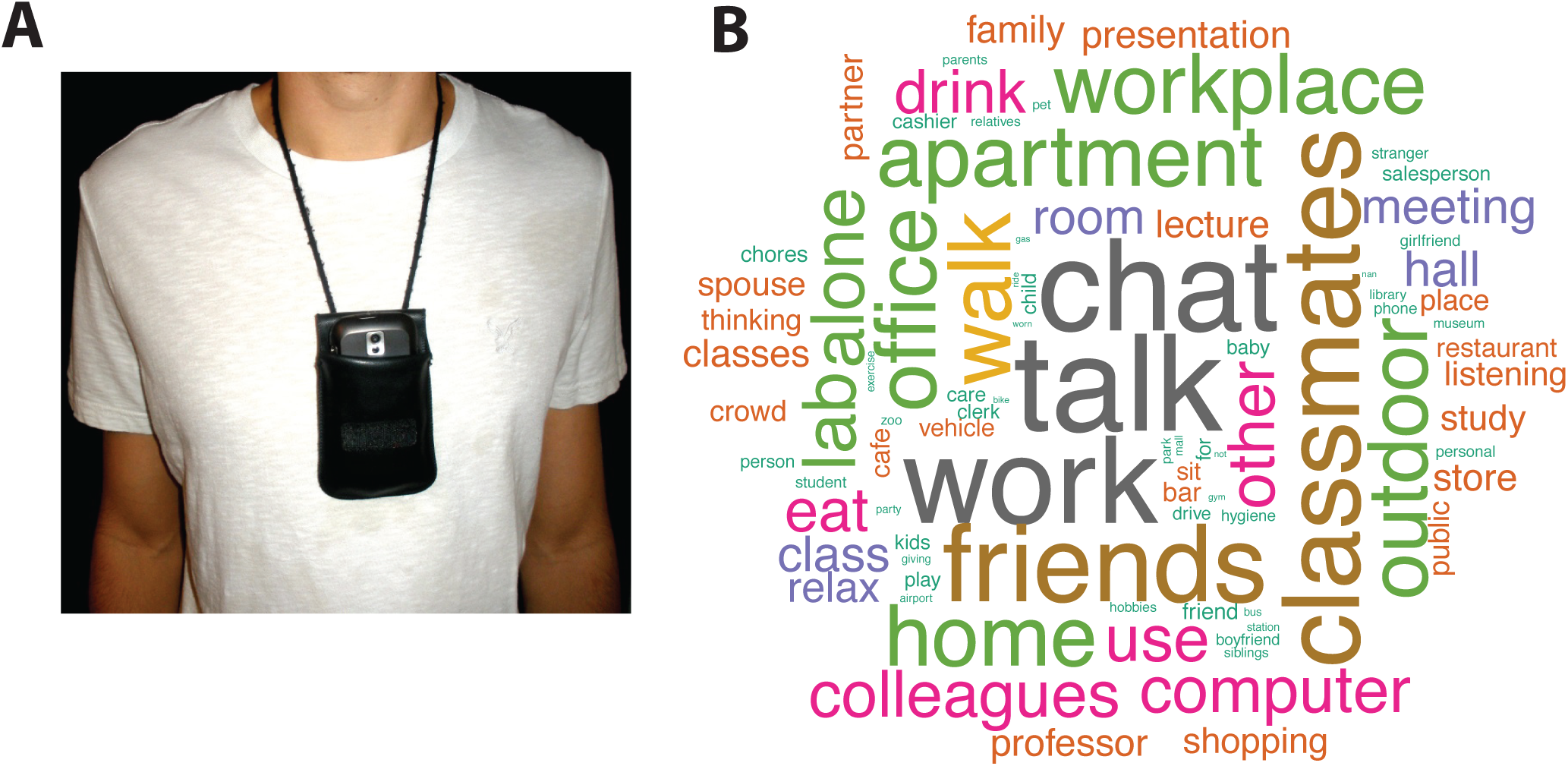
**(*A*)** The phone is worn around the neck with its camera exposed as shown. **(*B*)** A word cloud of the tags associated with the stimuli used in the fMRI experiment across all participants.

Two to four weeks after data collection, participants performed an autobiographical reminiscence task in an fMRI scanner where they were asked to relive experiences cued by images chosen from their own lives. We performed a Representational Similarity Analysis (RSA; Kriegeskorte et al., 2008) to identify the brain regions that represent personal semantics both generally and during vivid autobiographical reminiscence (see Materials and Methods and Figure 2).

**Figure 2.**
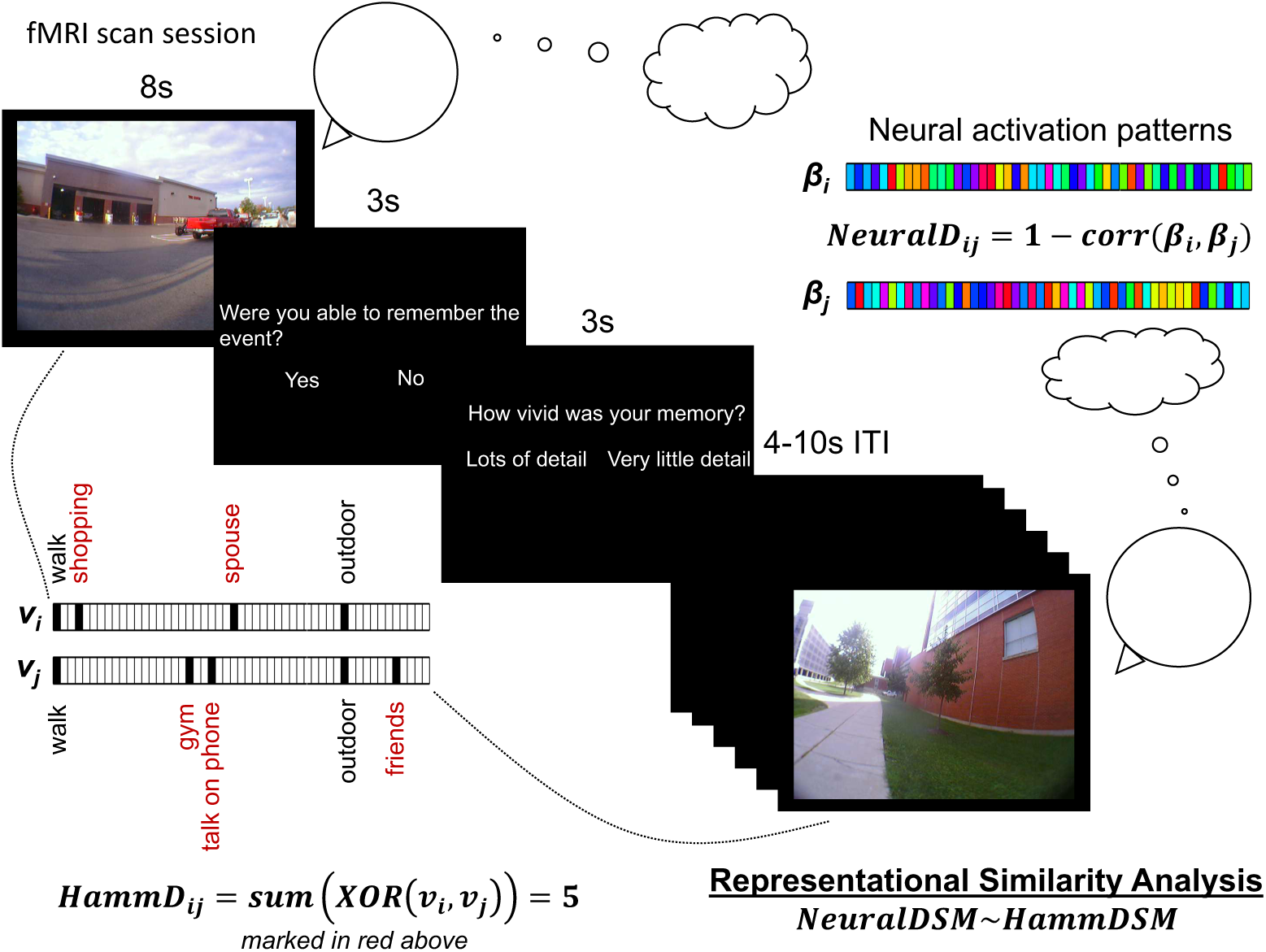
Depiction of the fMRI experiment and Representational Similarity Analysis (RSA). Participants are shown images from their own lives and are instructed to relive the associated experiences. The neural activity during this reminiscence period is analyzed using RSA to investigate whether distances between neural patterns (NeuralDij) corresponding to pairs of image cues (an example of such a pair is shown) relate to distances between the corresponding sets of semantic tags (HammDij). After the reminiscence period, participants indicate whether they remember the event and then report the vividness of their recollective experience.

### Behavioral Results

The Hamming distances between the tag sets for pairs of images across participants ranged from 0 to 15. 63.4 ± 4.7% SEM of the stimuli were reported as having produced successful reminiscence. The proportion of analyzed stimuli that were indicated as evoking vivid reminiscence by the nine participants ranged from 0.21 to 0.81 (mean 0.47 ± 0.07 SEM). The three different tag types were used to a similar extent across stimuli (Mean 94.9±1.5% SEM of the stimuli contained activity tags, Mean 91.2 ± 4.0% SEM contained people tags, and Mean 95.2 ± 1.3% SEM contained place tags). No differences were apparent in the percentage of stimuli that evoked vivid reminiscence depending on the type of tag present (Mean 44.5 ± 5.4% SEM of the stimuli with activity tags, Mean 42.4 ± 5.1% SEM of the stimuli with people tags, and Mean 44.3 ± 5.7% SEM of the stimuli with place tags evoked vivid reminiscence) suggesting that vividness was not linked to the presence of any particular tag type.

In order to understand the overall structure of events as organized by these tag sets across participants, we computed normalized pointwise mutual information (NPMI) between pairs of tags (see Materials and Methods for details). NPMI ranges from *-*1 to 1, with *-*1 indicating that the pair of tags never occurred together, 0 indicating that the tag occurrences were independent of each other, and 1 indicating that the tags co-occurred perfectly with each other. The NPMI matrix is presented in Figure 3A. We also plotted a network of tags with *N P M I >* 0.2 in Figure 3B to visualize the co-occurrence structures that emerge across participants. Together, the panels in Figure 3 demonstrate clusters surrounding university campus life and social/family life, reflecting general characteristics of the student pool from which we recruited participants.

**Figure 3.**
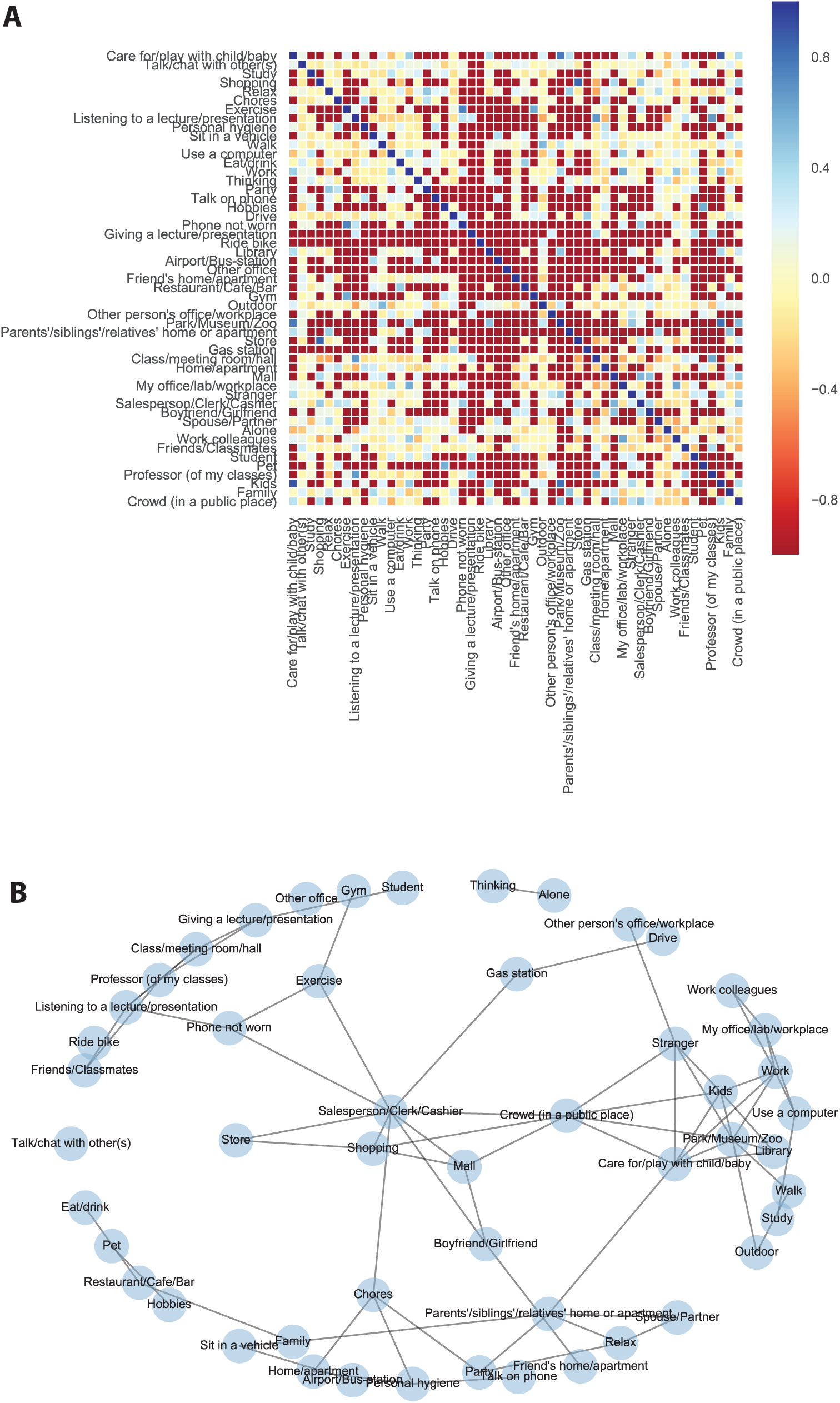
**(*A*)** Normalized pointwise mutual information (NPMI) between all pairs of tags, computed across participants. **(*B*)** A network of tags with *N P M I >* 0.2.

### Neural Results

RSA with the model in Equation 4 (a GLM relating neural distances with Hamming distances between tag sets) revealed a broad network of regions that represented personal semantics during the reminiscence task cued by participants’ own images. This general personal semantic network, shown in Figure 4A, included core parts of the default mode network (DMN) such as the precuneus, anterior cingulate, posterior cingulate, middle temporal gyrus bilaterally, and a right lateralized network including the medial and prefrontal cortices, parts of the inferior parietal lobule (supramarginal and angular gyri), and the parahippocampal cortex (see Supplementary Table S1 in Supplementary Section S1 for a complete list of regions with at least 10 voxels in the network). A personal image cue can trigger memory for general facts about similar events, which need not lead to detailed and vivid reminiscence of a specific event. Because Equation 4 did not include a term for vividness, *β*_*Hamm*_ tracks the regions that represent personal semantics generally and does not identify the regions that do so specifically during vivid reminiscence (implying greater episodic retrieval).

Therefore, we also performed an RSA with Equation 5 (a GLM relating neural distances with Hamming distances between pairs of tag sets, the overall level of vividness reported for the corresponding pairs of stimuli, as well as the interaction between vividness and Hamming distance) in order to identify both the network that represented personal semantics for vividly re-experienced events (i.e., the main effect of *Hamm*) and the regions that represented the subjective contents of experience for vivid but critically not for non-vivid memories (i.e., the conjunction of *Hamm* and *-Hamm * Vivid*). Figure 4B shows the regions that represent personal semantic content during vivid reminiscence. This is mostly a sub-network of the general personal semantics network, but relatively more right lateralized and therefore this “vivid”personal semantic network also includes parts of the DMN, such as the precuneus bilaterally, and in the right hemisphere, posterior cingulate, parahippocampal cortex, medial and pre-frontal cortices (see Supplementary Table S2 in Supplementary Section S2 for a complete list of regions with at least 10 voxels in the vivid personal semantics network).

**Figure 4.**
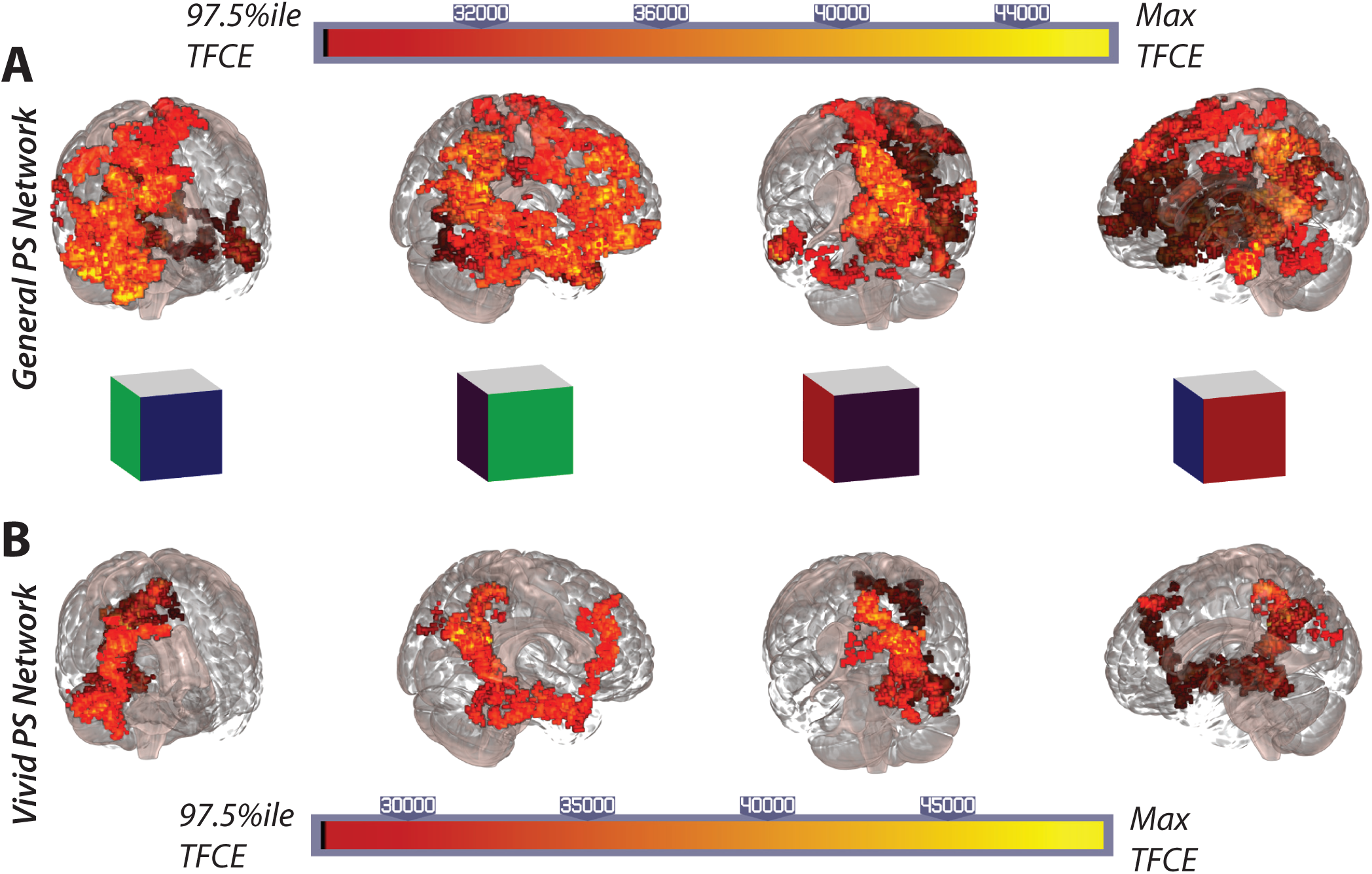
**(*A*)** The network of regions involved in the representation of general personal semantics as identified by the RSA analysis in Equation 4. Four different views of a glass brain are shown. From left to right, the blue face of the orientation cube corresponds to the front of the brain, the green face stands for the right hemisphere, the black face for the back, and the red face for the left hemisphere. **(*B*)** The network of regions involved in the representation of personal semantics during vivid reminiscence as identified by the RSA analysis in Equation 5. The same views presented in **(*A*)** are shown and comparing the two networks reveals that the vivid reminiscence network is a subset of the more general personal semantics network identified in **(*A*)**.

While the *Hamm* term in Equation 5 tracks the regions involved in representing personal semantic content during vivid reminiscence, it does not address whether those regions distinguish between vivid and non-vivid recollection. This distinction between vivid and non-vivid reminiscence is captured by the conjunction between *Hamm* and*-Hamm * Vivid* which identifies regions where *Hamm* predicts neural distances for vivid pairs of images but does so to a significantly less extent for non-vivid pairs (see Materials and Methods). A dominant cluster in the right precuneus (Figure 5) is identified as the region representing self-relevant contents of an experience when vivid autobiographical memory is generated but critically, the right precuneus content representations are significantly attenuated when the memory is non-vivid (see Table 1 for the MNI coordinates of the peak voxels in regions with at least 10 voxels in the vivid-only personal semantics network).

**Table 1.** Peak voxel coordinates of regions with at least 10 voxels in the vivid-only personal semantic network(Figure 5). The FSL-Harvard-Oxford cortical-subcortical atlas was used to get coordinates in MNI space. When multiple sets of coordinates are shown for a region, they correspond to multiple peak voxels.

| Region | Voxel count | Mean TFCE | Max TFCE<sup>1</sup> | MNI coordinates |  |  |
| --- | --- | --- | --- | --- | --- | --- |
|  |  |  |  | *x* | *y* | *z* |
| R. Precuneus | 135 | 17412.7 | 19694.3 | 17.5 | -65.5 | 23 |
| R. Cuneal Cortex | 31 | 17887.6 | 19738.4 | 17.5 | -68 | 23 |

<sup>1</sup> 95th percentile TFCE threshold = 16232.6, Max network TFCE = 19738.4

**Figure 5.**
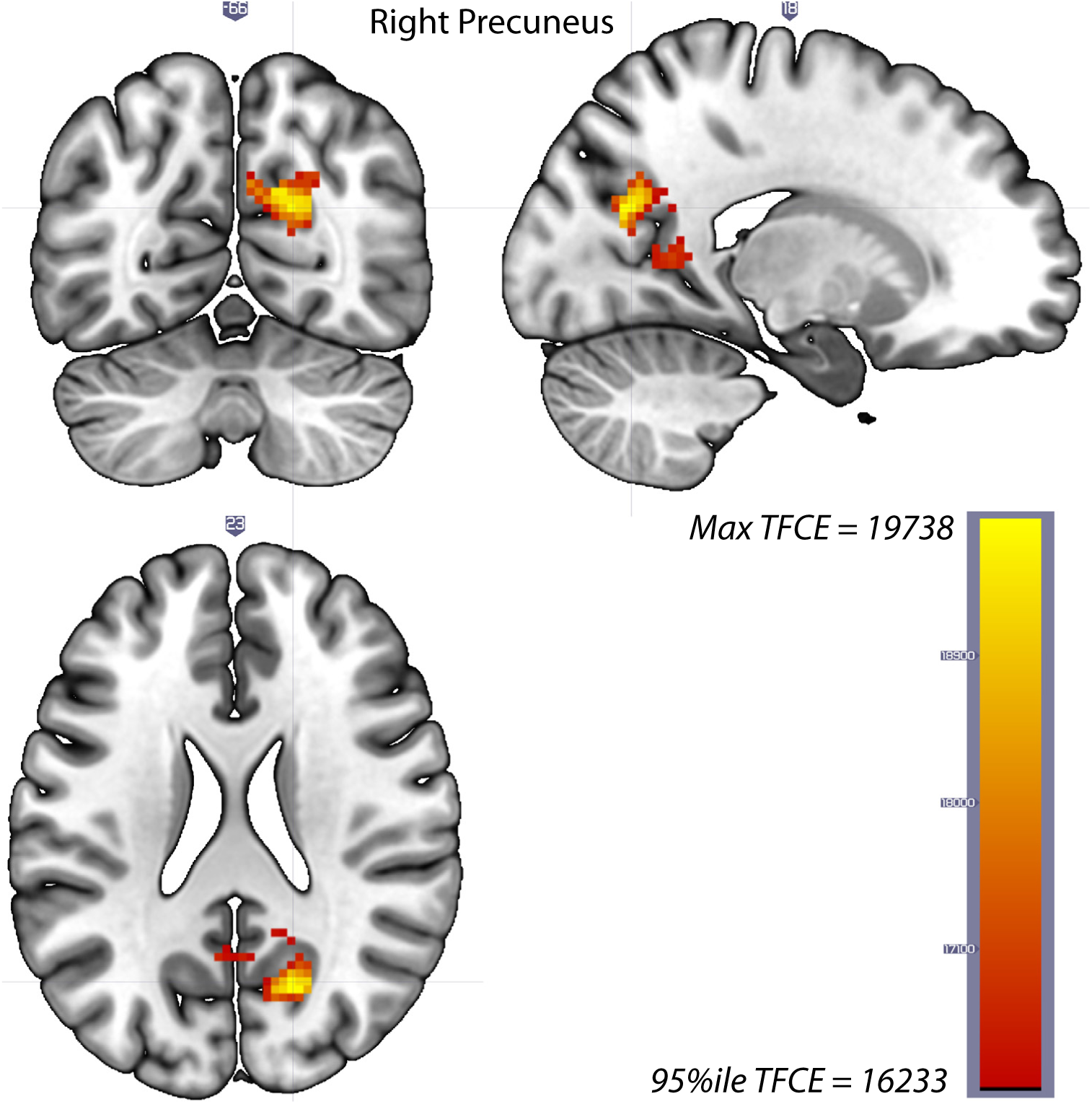
The right precuneus represents personal semantics during vivid reminiscence but to a lesser extent during non-vivid reminiscence (conjunction analysis based on Equation 5, Materials and Methods)

Finally, we present partial residual plots to visualize the relationship between *Hamm* and neural distances in the right precuneus after taking into account the contribution from the other independent variables in Equation 5 (Materials and Methods), and we do this separately for vivid and non-vivid pairs. Since overlaying the residuals obscures the differences in the slopes of the regression lines between vivid and non-vivid conditions, we opted to display only the regression lines in Figure 6 and the individual participants’ plots with partial residuals overlaid in Supplementary Section S5, Figure S2. Figure 6A shows that neural distances in a sphere surrounding the peak right precuneus voxel are related to Hamming distances between the tag sets of vivid pairs of stimuli (*Vivid* >0 in Equation 5) and Figure 6B demonstrates that this relationship is considerably attenuated for non-vivid pairs of stimuli (*Vivid* ≠ 0 in Equation 5). These differences in how neural distances relate to dissimilarities between personal semantic tags between vivid and non-vivid pairs suggest that vivid reminiscence is accompanied by activity in the right precuneus reflecting higher fidelity self-relevant personal semantic representations relative to non-vivid reminiscence.

**Figure 6.**
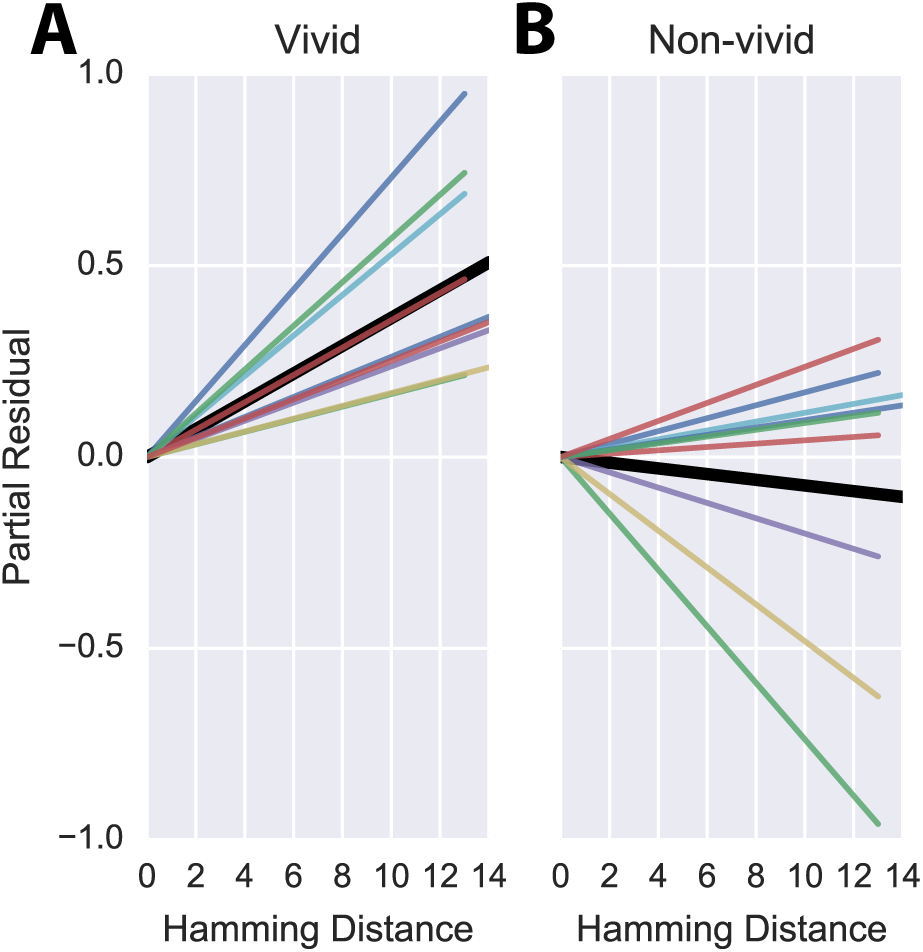
The slopes of the regression lines in Equation 5 describing the relationship between neural distances and Hamming distances between the tag sets in a sphere of radius 7.5 mm around the peak voxel in the right precuneus. **(*A*)** The colored lines show individual participants’ regression lines for the relationship between Hamming distance and neural distance for vividly remembered pairs of images after accounting for the contribution from other independent variables in Equation 5 (i.e., the partial residual). The slope of the solid black line is the mean over the individual regression lines. **(*B*)** The relationship between Hamming distance and neural distance for the less vividly remembered pairs of images after accounting for the contribution from other independent variables in Equation 5.

## Discussion

In a recent review, Renoult et al. (2012) identified the neural correlates of personal semantics, thought to consist of facts about one’s own life extracted over many repeated experiences. Renoult et al. (2012) described a personal semantics network that included the mPFC, retrosplenial cortex, temporal pole, posterior temporal cortex, precuneus, middle and inferior temporal gyri, inferior parietal lobe, hippocampus, parahippocampal gyrus, temporo-parietal junction, ventrolateral prefrontal cortex, and fusiform gyrus. The specific neural correlates depended on where the specific operationalization of personal semantics was located in the spectrum from semantic to episodic memory. The personal semantics network we identified in an autobiographical reminiscence task (Figure 4A, Supplementary Table S1) overlaps highly with the broad network described in Renoult et al. (2012) and includes core parts of the default mode network (DMN), which is thought to be involved in the processing of self-relevant information and in unconstrained mind-wandering. The DMN overlaps highly with contextual association networks and Bar et al. (2007) suggested that unconstrained thought processes, much like explicit associative memory processing, involve activation of such associations. Therefore, it is perhaps unsurprising that the network involved in instantiating associated personal semantic representations upon viewing an autobiographical image-cue is congruent with the associative-default network (see Figure 1 in Bar, 2007).

We also identified a network that represented personal semantic content for vivid memories. This set of regions (Figure 4B, Supplementary Table S2) was mostly a sub-network of the more general semantic network but relatively more right lateralized. The posterior parietal cortex (PPC; including the posterior cingulate and precuneus) is a dominant part of the retrieved personal semantics networks we identified. Though studies of the human PPC have traditionally focused on visuospatial and sensorimotor functions, the PPC has received increased attention recently as a region that plays an important role in episodic memory (for reviews, see Cabeza et al., 2008; Ranganath and Ritchey, 2012; Sestieri et al., 2017; Vilberg and Rugg, 2008; Wagner et al., 2005). Though previous studies showed a predominantly left lateralized parietal retrieval network, Vilberg and Rugg (2008) suggested that it could have been a result of the limited range of materials (mostly verbal) used in those studies (e.g. see Simons et al., 2008 for evidence that source recollection of faces vs words evokes more activity in the right hemisphere, but see Duarte et al., 2011 for an argument that retrieval MTL and PPC networks are material-general). Therefore, our observation of a right lateralized personal semantic network associated with vivid reminiscence could be explained by our use of highly personally relevant image cues drawn from participants’ own lives. However, since all the participants in our study were female, we are unable to rule out an alternative gender-based explanation for the right lateralization (but Viard et al. (2007), with all twelve participants being female, reported a left-lateralized network in an autobiographical memory retrieval task that used verbal cues collected from family members prior to fMRI scanning).

Finally, given that vividness is a defining feature of successful autobiographical recollection, we focused on the regions within the broader network that represented retrieved personal semantic content specifically in service of vivid reminiscence, but not during non-vivid recall. The conjunction analysis identified the right precuneus (pC) as the locus of representation of content specifically accompanied by vivid reminiscence, but, critically, personal semantic representations were significantly attenuated in the right pC during non-vivid relative to vivid recall. Both univariate and multivariate activity in the pC is consistently related to vividness ratings across AM experiments (Bird et al., 2015; Cavanna and Trimble, 2006; Gilboa et al., 2004; Richter et al., 2016; St-Laurent et al., 2015). Furthermore, Gilboa et al. (2004) presented family photographs, which are closer to the type of stimuli we used and they found that univariate activity in the right pC and bilateral lingual gyri was associated with vividness ratings. They suggested that vivid and detailed AM was required to engage the posteromedial cortex, which is thought to represent contextual details (*cf.* Ranganath and Ritchey, 2012). Our results offer direct evidence for this idea by demonstrating that neural activity patterns during vivid but not during non-vivid recall in the right pC represent the specific self-relevant contents of the original experience as indicated by participants.

The precuneus has been called the “mind’s eye”(Fletcher et al., 1995) and pC activity is consistently associated with mental imagery and episodic memory (Cabeza and Nyberg, 2000; Cabeza and St. Jacques, 2007). A special status for the pC has been proposed within the default mode network (Buckner et al., 2008; Cavanna and Trimble, 2006; Leech et al., 2012; Utevsky et al., 2014). Fransson and Marrelec (2008) performed a partial correlation-based connectivity analysis which measured the extent of interaction between nine nodes within the DMN after subtracting out the common influences from other nodes. This was done for resting state as well as a working memory task and they showed that pC was the only node that exhibited strong connectivity with virtually every other node. Functional connectivity analysis (Baird et al., 2013) and anatomical coupling and voxel-based morphometry analyses (McCurdy et al., 2013) have suggested an important role for the pC in metacognitive ability for memory retrieval. These connectivity patterns taken together with our results suggest that the pC may play an important role in the integration of personal semantic information from other parts of the network leading to a detailed representation of the self-relevant contents of a specific experience, supporting the subjective experience of vivid autobiographical reminiscence.

The idea that the pC may have a privileged status within the DMN is further supported by the discovery that along with regions in the MTL, the pC is one of the first regions to be affected in early Alzheimer’s disease (AD) (Jack et al., 2009). There is catastrophic breakdown of information flow when a hub in a network is affected (Albert et al., 2000). This could explain why in the early stages of AD, people lose track of time, people, and places (also see Peer et al., 2015 for evidence that the same regions are important for mental orientation along the different dimensions of space, time, and persons and that the pC activated across these domains). On a related note, a new memory syndrome, severely deficient autobiographical memory (SDAM), was identified recently (Palombo et al., 2015) in three healthy adults with otherwise normal cognitive functioning who were severely impaired on autobiographical memory function. This impairment was specific to vivid visual episodic re-experiencing of personal events but did not extend to remembering personal semantics. Furthermore, even though they were impaired relative to the controls in reporting spatiotemporally specific episodic details of remote events, they were able to produce episodic details for recent events, albeit accompanied by significantly reduced vividness ratings across all time periods. fMRI scans during a cued-autobiographical recall task revealed that there was reduced activity compared to the controls in areas including the left mPFC and right precuneus. Our results are consistent with Palombo et al. (2015)’s report and suggest that the subjective experience of vivid reminiscence is facilitated by activity in the right precuneus reflecting personal semantics of retrieved episodic details whereas personal semantics more generally are represented by a broader network of regions, which can explain the selective vividness deficits but intact personal semantics in people with SDAM.

## Conclusion

It has been suggested that AM retrieval is guided by semantic retrieval (cf. Conway and Pleydell-Pearce, 2000). We identified the general network, including core parts of the default mode network, that represents retrieved personal semantics during AM search over several weeks of real-world experience. The precuneus is a hub within this network (Cavanna and Trimble, 2006; Damasio, 1989; Fransson and Marrelec, 2008; also see Binder and Desai, 2011; Moran et al., 2013) and our results suggest that activity in the precuneus supports the subjective experience of vivid reminiscence by representing personal semantic attributes with higher fidelity during vivid compared to non-vivid recall. This account provides a plausible mechanism by which people make metacognitive judgments about their recollective experiences, and may provide key support to theories that suggest a critical role of the precuneus in the autobiograpical memory deficits seen in Alzheimer’s disease and other forms of dementia.

## Materials and Methods

### Device and Software

Each participant carried an Android-based smartphone in a pouch attached to a neck strap as shown in Figure 1A from morning until evening. The smartphone was equipped with a custom lifelogging application that acquired image, time, audio (obfuscated), GPS, accelerometer, and orientation information throughout the day and uploaded those data to a secure remote server when the smartphone was connected to a charger and detected WiFi. This transmission usually happened once per day at the end of the day because users charged the phone overnight. The data were sent in batch mode via SFTP (Secure File Transfer Protocol) for added security and remained inaccessible to other users in the system. The participants had control over what data they wanted to share with the experimenters. They were instructed on how to delete data from the phone and from the server. They were also allowed to turn the application off or to place a flap over the camera lens at any time during the data collection period when they felt the need for privacy. The lifelogging application was written by our programmers using Java (Oracle Corporation) to run in the background as a service. Data acquisition times could be fixed or variable, and they were determined by a movement based trigger to preserve battery resources when the user was not very active.

### Participants

Participants were recruited using advertisements placed on notice boards in multiple buildings on the main campus of The Ohio State University. To join the study, potential participants had to be willing to participate in the lifelogging data collection and to be willing and able to undergo an MRI scan. They were compensated at the rate of $10 per day for wearing the smartphone to collect data and at the rate of $15 per hour for the fMRI session. We recruited 10 participants (aged 19–26 y, mean age = 21.4 y; nine female), nine of whom wore the smartphone for *∼* 1 month. The tenth participant wore the smartphone for 2 weeks. One participant (male) did not complete the fMRI session due to discomfort in the scanner; therefore, we did not include the data for that participant in any of our analyses. Our study has a similar number of participants as other fMRI studies using lifelogging devices (13 participants and 10 days of lifelogging in Cabeza and St. Jacques, 2007; 10 participants and 2 days of lifelogging and a 5 month follow-up in Milton et al., 2011).

### Ethics Statement

The research protocol was reviewed and approved by the Institutional Review Board at The Ohio State University. Written informed consent was obtained from all participants, once before the lifelogging data collection phase and once before the fMRI session.

### Behavioral Tasks

There were two main behavioral tasks that were performed before the MRI session. The first behavioral task was performed each evening during the lifelogging period. After the smartphone was connected to a power outlet to be charged overnight and had uploaded the data to our server, participants reviewed the images from that day through a web interface, a link to which was uniquely generated for each participant and provided to the participant before data collection, segmenting their stream of images into distinct episodes and tagging each episode with a set of tags chosen from a drop-down menu (Table 2). Participants were instructed to choose tags that best captured the contents of that episode and those that were likely to be good memory cues. The tags belonged to one of three categories: places, activities, and people. If no tag fit the episode, participants could choose “other”. For each episode, they also provided a brief title and description. Insofar as only the participant knew the right tag to pick for a given episode, the set of tags captures the subjective contents of that episode. For instance, looking at someone else’s data with images of a person in it, it may be difficult to pick the appropriate tag from amongst “Spouse/Partner”, “Boyfriend/Girlfriend”, “Family”, “Work colleagues”, “Stranger”, and “Friends/Classmates”. While other tags are more objective, such as “Salesperson/Clerk/Cashier”or “Gas station” the chosen tags are nevertheless the aspects chosen by the participant as the most salient of that episode from potentially many other descriptors. Therefore, the current analyses which are based on participant-generated content tags capture more self-relevant and subjective aspects of experience than did our previous work (Nielson et al., 2015) which based on objective GPS locations and timestamps. A word cloud of the tags belonging to the episodes used in the fMRI experiment across all nine participants is shown in Figure 1B. The second behavioral task was conducted midway through the lifelogging period and at the end of the lifelogging period. After they collected data for two (and/or four) weeks, participants came into the laboratory on the Thursday of the third (and/or fifth) week and were tested over their ability to identify when events depicted in images drawn from his/her own lifelogs occurred. Specifically, they were shown a series of images from the weekdays of the preceding 2 weeks on the computer screen one at a time and asked to determine whether the image was from the first week or the second week. The results of this week discrimination task will be reported in a separate paper.

**Table 2.** The 51 tags available to participants across three categories: places, activities, and people. The number of available tags in each category are in brackets. Additionally, they could also choose “other”if none of these fit the event.

| Category | Tags |
| --- | --- |
| Places (16) | Outdoor, Airport/Bus-station, Gas station, Park/Museum/Zoo, Gym, Library, Parents’/siblings’/relatives’ home or apartment, Mall, Friend’s home/apartment, Class/meeting room/hall, Restaurant/Cafe/Bar, My office/lab/workplace, Home/apartment, Other office, Store, Other person’s office/workplace |
| Activities (22) | Chores, Thinking, Party, Talk on phone, Use a computer, Exercise, Shopping, Personal hygiene, Relax, Eat/drink, Talk/chat with other(s), Phone not worn, Study, Work, Drive, Care for/play with child/baby, Ride bike, Giving a lecture/presentation, Listening to a lecture/presentation, Walk, Sit in a vehicle, Hobbies |
| People (13) | Kids, Family, Friends/Classmates, Pet, Salesperson/Clerk/Cashier, Boyfriend/Girlfriend, Stranger, Alone, Professor (of my classes), Student, Spouse/Partner, Crowd (in a public place), Work colleagues |

### Analysis of tag co-occurrence structure

In order to characterize the co-occurrence structure of semantic tags that emerges across participants, we computed pointwise mutual information (PMI), a measure of association between two features. PMI for a pair of tags *x* and *y* is given by:

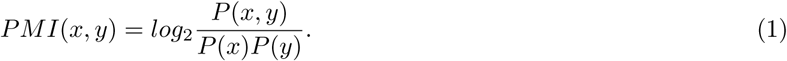

The probabilities in Equation 1 are calculated by accumulating frequencies of tags as well as frequencies of co-occurrences of tag pairs in all events across participants and then dividing by the total number of events (120 × 9 = 1080). PMI is sensitive to tag frequency and is bounded between *-∞* and min[*-log*_2_*p*(*x*)*, -log*_2_*p*(*y*)]. Therefore, we used the normalized pointwise mutual information (NPMI) which is more easily interpretable and is less sensitive to tag frequency:

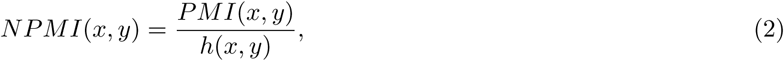

*Where*

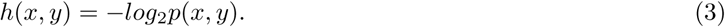

*NPMI*(*x, y*) = *-*1 when the pair of tags never co-occurs, *N P M I*(*x, y*) = 0 indicates that the tag occurrences are independent of each other, and *N P M I*(*x, y*) = 1 indicates that the tags always co-occur.

## MRI Acquisition

MRI data were acquired on a 3-T Siemens Magnetom Trio TIM system with a 16-channel head coil. Anatomical images were acquired with a sagittal, T1-weighted, magnetization prepared rapid acquisition gradient echo sequence [1.0-mm isotropic voxels, repetition time (TR) = 1900 ms, echo time (TE) = 4.68 ms, 160 slices with field of view (FoV) = 256 mm]. Functional images were acquired with an echoplanar imaging sequence (2.5-mm isotropic voxels, TR = 3000 ms, TE = 28 ms, flip angle = 80°, 47 slices with FoV = 250 mm).

## Stimuli Selection

We selected 120 images from each subject’s lifelogging data to present to the subject in the scanner. First, we excluded pictures of floors/ceilings/walls, blurry images, and images with inadequate exposure. Then, we selected images that appeared to have enough detail that they could act as cues for distinct episodes. From this subset of images, we selected images representing events that spanned the entire period each participant wore the lifelogging device, with as uniform sampling of events as possible.

**fMRI Experiment**

In the scanner, participants were instructed that they would be viewing images from the experience sampling experiment they recently completed and told that each image would be displayed for 8 s. Participants were asked to “…try to remember the event depicted in the picture, and try to relive your experience mentally.”After the remembrance period for each event, participants were asked if they remembered the event (“yes”or “no”) and how vividly they recalled the event (“lots of detail”or “very little detail”). Participants were given 2.5 s to respond to each of those questions using a button box held in their right hand. The images were presented in random order, and the task was split into eight runs with 15 images in each run. With each image presented for 8 s and each question for presented 2.5 s with a 0.5 s interstimulus interval, each trial took a total of 14 s. The intertrial interval was jittered uniformly between 4 and 10 s, allowing for a true event-related design.

## fMRI Processing

fMRI processing was carried out using Analysis of Functional NeuroImages (AFNI) (Cox, 1996) and Functional Magnetic Resonance Imaging of the Brain (FMRIB) Software Library (FSL) (Smith et al., 2004). The T1-weighted anatomical image was intensity-normalized, skull-stripped, and warped to a 2.5-mm MNI-152 template using *3dQwarp*. We selected a 2.5 mm template to match the resolution of the functional scans. For the functional scans, we dropped the first two TRs of each run, then removed spikes with *3ddespike* and temporally shifted all of the slices in each volume to the start of the TR using *3dTshift* with Fourier interpolation. We then warped the functional scans to template space, blurred them to 4 mm FWHM using *3dBlurtoFWHM*, and scaled the voxel values to a mean of 100 (maximum of 200) for each run. At this point, we performed independent component analysis of each functional run with FSL’s *MELODIC*. Components were visually inspected to identify noise components following published guidelines (Kelly et al., 2010). Noise components were regressed out of the functional runs using FSL’s *fsl_regfilt* command. We then ran a regression with restricted maximum likelihood estimation of temporal autocorrelation structure on the filtered functional runs using *3dDeconvolve* and *3dREMLfit* to generate single-trial betas for each reminiscence trial and to regress out the effects of the mean and derivative of motion terms, as well as cerebrospinal fluid signal. The regressor for each image presentation was an 8-s block convolved with a hemodynamic response function. The neural activity of the question prompts were accounted for with a 2.5 s block convolved with a hemo-dynamic response function. We modeled response processing and motor activity related to the button push with a set of nine tent functions over the 16 s after the question response. Including these tent functions in our model allowed us to estimate the motor response robustly for each subject so that the signal from the motor responses did not contaminate the single-trial beta fit for each reminiscence period. Lastly, we regressed out local white matter signal with *3dAnaticor*. Researchers were not blinded during preprocessing or subsequent analyses.

## Representational Similarity Analysis

Representational Similarity Analysis (RSA, Kriegeskorte et al., 2008) is a data-analytic framework that allows us to quantify the relationship between the multivoxel patterns of neural activity and the behavior of interest. We used RSA to predict dissimilarities between the neural representations of events based on the dissimilarities between the events in terms of their subjective contents as captured by the tags provided by participants during the lifelogging phase as well as the vividness ratings provided during the reminiscence task in the scanner. See Figure 2 for a depiction of the task and analysis.

For each pair of images presented to the participants, we calculated the Hamming distance between the associated tag sets. Since a total of 52 unique tags were used (including the “other” tag), each tag set can be represented as a 52-dimensional binary vector where each entry denotes the presence/absence of a tag. The Hamming distance between two binary vectors A and B is simply the number of positions where they differ, or in other words, Hamming distance = sum(XOR(A,B)). For example, if A = [1 1 0 0 1 0 1…] and B = [0 0 1 1 0 0 1…] with only the first 5 positions being different, the Hamming distance is 5. As a more concrete example, if image A had been tagged with Walk, Outdoor, Talk on phone and image B had the tags Walk, Store, Talk on phone, the Hamming distance between them is 2 since there are 2 tags that are different between the two sets reflecting the difference in location between the two otherwise similar events.

In our previous analysis (Nielson et al., 2015), for each pair of images presented to the participants, we calculated the geodesic distance in meters between the two GPS coordinates and the difference in time in seconds. Geodesic distance was calculated using the GeoPy Python package. Image pairs with spatial distances less than 100 m were excluded because these distances are below the reliability of the GPS radios in these smartphones. Image pairs with temporal distances below 15.6 h were excluded based on prior work because of a discontinuity in the spatiotemporal distribution of image pairs (Sreekumar et al., 2014). The discontinuity between 14 and 16-h results from participants taking off their cameras to sleep. This gap is propagated through the rest of the results as a relative lack of image pairs that are multiples of *∼* 15 h apart. An analysis of the structure of similar lifelogged images demonstrated that image pairs taken from identical spatiotemporal locations occupied a lower dimensional manifold than those image pairs taken from separate spatiotemporal locations (Sreekumar et al., 2014;2017). By removing image pairs separated by less than 100 m and 15.6 h, we reduced the possibility that the images themselves would give rise to the present results as a consequence of within- and between-episode image properties. Some participants spent time out of town during the period of data collection, resulting in a small portion of image pairs with spatial distances greater than 30 km; these image pairs were also excluded in Nielson et al. (2015) and we impose the same spatial limit. Images that were blurry or contained reflections of the participants were also excluded. Hamming distances between the tag sets of the remaining pairs of image stimuli were calculated as described earlier. In order to further control for visual similarity, we compared five different popular image representations based on how well they identified temporally close images as visually similar and chose the color correlogram representation (Huang et al., 1997; Supplementary Section S3 and Figure S1 for details). Euclidean distances between the correlogram image representations were computed and entered into the General Linear Models (GLMs) described below as a visual control (*V isSim* inEquation 4 and Equation 5).

In order to investigate both cortical and sub-cortical contributions to content retrieval, we performed a whole-brain searchlight analysis (Kriegeskorte et al., 2006) using the PyMVPA package (Hanke et al., 2009). Representational similarity analysis (RSA) was performed on voxels within spherical neighborhoods of 7.5 mm radius surrounding a central voxel. An initial 2.5 mm resolution gray matter mask in the Montreal Neurological Institute (MNI-152) standard space was used to input the fMRI data to the searchlight function but for each individual, we used a subject-level gray matter mask warped to MNI-152 space to select the spheres on which to run the analysis. Within each sphere, the neural distance for each image pair was calculated as 1 minus the Pearson correlation between the voxel-level single-trial betas for the trials corresponding to those image pairs. Neural distances were z-scored within participants. In each searchlight sphere in each subject, we ran the following GLM:

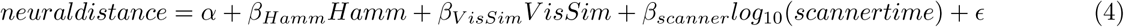

Scanner time was calculated as the number of seconds between presentation of the images during the fMRI experiment. We used the log of time based on previous literature that has shown a power-law relationship for neural representations (Gallistel and Gibbon, 2000). In each sphere, we performed a t-test on the betas from the subject-level GLMs to determine if they were significantly different from zero across participants. We used nonparametric permutation to test for significance (Ernst, 2004) because the pairwise nature of the distances in our analysis violated the assumption of independent samples. Neural data were permuted with respect to behavioral data within participants. This process was repeated for 1000 permutations of the neural data. We performed threshold-free cluster enhancement (TFCE; Smith and Nichols, 2009) on the Hamm t-value maps for both the unpermuted data as well as for the 1000 permutations. The maximum and minimum TFCE values across all spheres for each permutation were recorded. The 97.5th percentile of the max TFCE values was chosen as the threshold above which a positive TFCE value in the unpermuted data is deemed to be significant. Similarly, we tested the negative end by using the 2.5th percentile of the min TFCE values as the threshold (this procedure is essentially a two-tailed test at p= 0.05). This analysis reveals the clusters of brain regions whose activity patterns reflect the relationships (captured by Hamming distances) between events in terms of their contents. Additionally, we wanted to identify regions that may support metacognitive judgments (such as vividness of the recollective experience) based on the contents of memory retrieval. One possibility is that the quality of personal semantic representations in such brain regions would differ between different levels of reported vividness. Therefore, we ran the following model to investigate the brain regions that represent subjective content during vivid but not during non-vivid reminiscence:

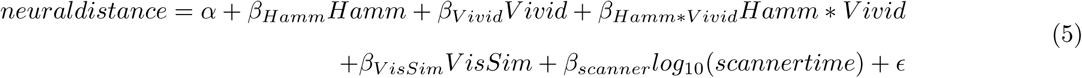

For a given pair of stimuli, *Vivid* was coded as 0 if both were reported to be vividly recalled, 0.5 if one of them was vivid, and 1 if neither was vivid. After subtracting out the effects of temporal proximity and visual similarity of the stimuli in the scanner, this coding scheme allows us to interpret *β*_*Hamm*_ as describing the relationship between Hamming distances and neural distances for vividly remembered events since the other terms vanish for *Vivid* ≠ 0. We expected the interaction between Hamming distance and vividness to be negative as that would indicate that the effect of Hamming distance is greater for vividly remembered events relative to less vividly remembered events in its ability to predict the neural distances between them. To identify the regions that show a significant effect of both Hamming distance by itself as well as the negative interaction with vividness, we performed a conjunction analysis by taking *M in*(*t*_*Hamm*_*, -t*_*Hamm*V* *ivid*_). TFCE was performed on this minimum t-statistic map and the permutation procedure was performed as earlier to assess significance of the clusters. 95th percentile of max TFCE across permutations was used to test significance since this was a one-tailed directional test. The conjunction analysis reveals the regions that reinstate subjective contextual details to a greater extent for vividly remembered events relative to the less vivid or non-vivid events.

Finally, to visualize the relationship between Hamming distances and neural distances in a sphere (radius = 7.5 mm) surrounding the peak voxel in the right precuneus, we used partial residual plots. Partial residual plots describe the relationship between a dependent variable and an independent variable after accounting for the contribution from other independent variables in a multivariate regression model. Specifically, to visualize the relationship between Hamming and neural distances for vividly remembered pairs of images (*Vivid* = 0 in Equation 5), we first computed residuals by regressing neural distances versus all the independent variables in Equation 5) for vivid pairs. *β*_*Hamm*_*Hamm* is then added to these residuals to get partial residuals = residuals + *β*_*Hamm*_. The partial residuals are plotted against *Hamm* to visualize the relationship between *Hamm* and neural distances for vivid pairs after taking into account the effect of all the other independent variables. This procedure can be understood intuitively if one considers the hypothetical case when all the other independent variables explain the response variable perfectly.In that case, residuals = *-β*_*Hamm*_*Hamm* and therefore the partial residuals after adding *β*_*Hamm*_*Hamm* back in would be 0. The regression lines overlaid on the partial residuals vs Hamming distance plot have the same slope as in the full model (i.e., *β*_*Hamm*_) but have an intercept of 0. Similarly, we plot the partial residual plot for the less vivid and non-vivid (*Vivid* ≠ 0) pairs, but now for components *β*_*Hamm*_*Hamm* + *β*_*Hamm*V* *ivid*_*Hamm * Vivid* since the relationship between Hamming distances and neural distances now also depends on *Vivid* via the interaction term (which was 0 for vivid pairs).

## Acknowledgements

This work was partially supported by Air Force Office of Scientific Research (AFOSR) grant FA9550-09-1-0614 (SD), National Science Foundation (NSF) grant 1631403 (PBS), and a grant from the Rudi Schulte Research Institute (PBS). We would like to thank Ben Stone and Ben Wierwille for their work on the lifelogging app. This work was supported in part by an allocation of computing time from the Ohio Supercomputer Center.

## Supplementary Information

### S1 List of regions in the general personal semantic network

**Table S1.**
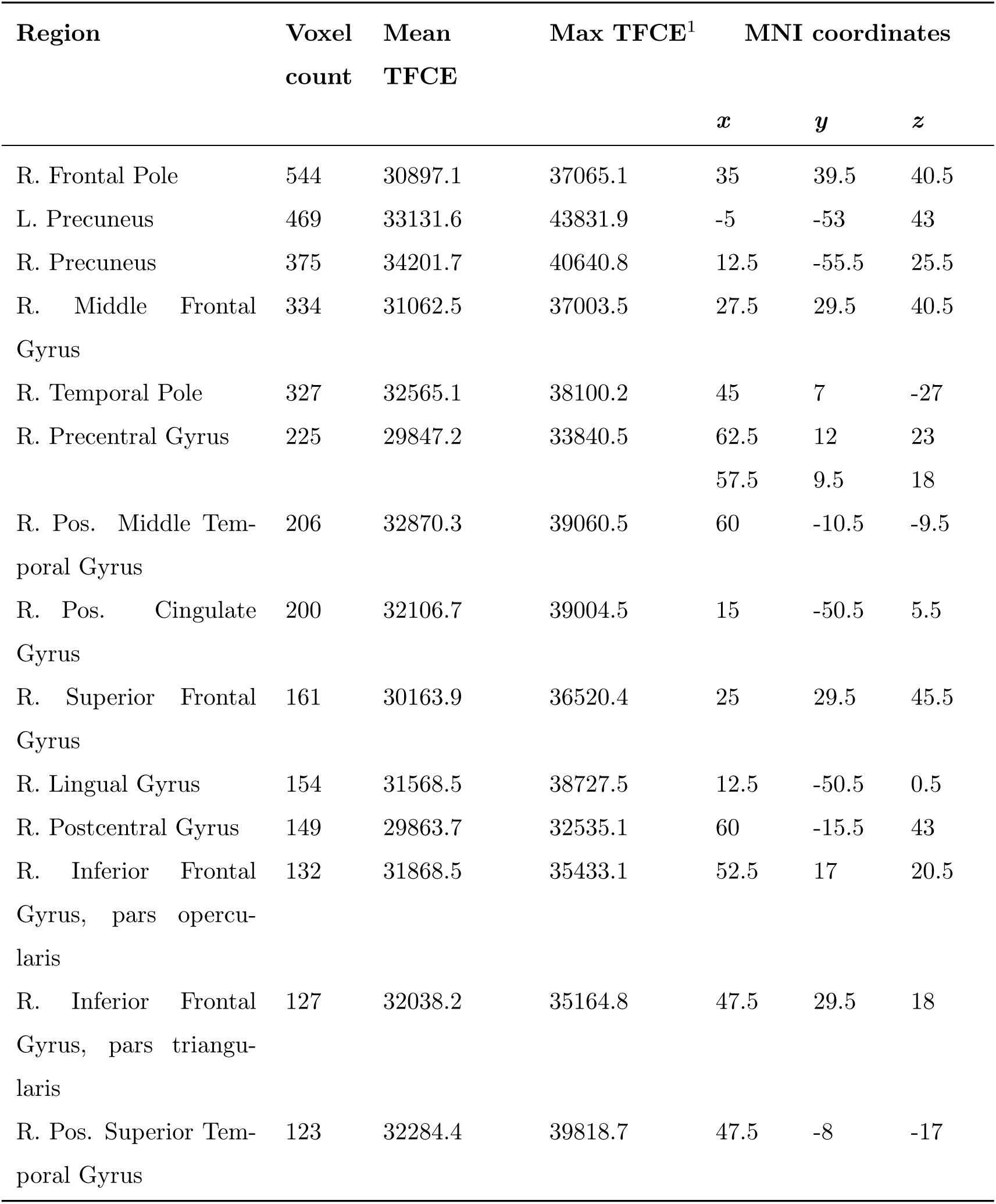
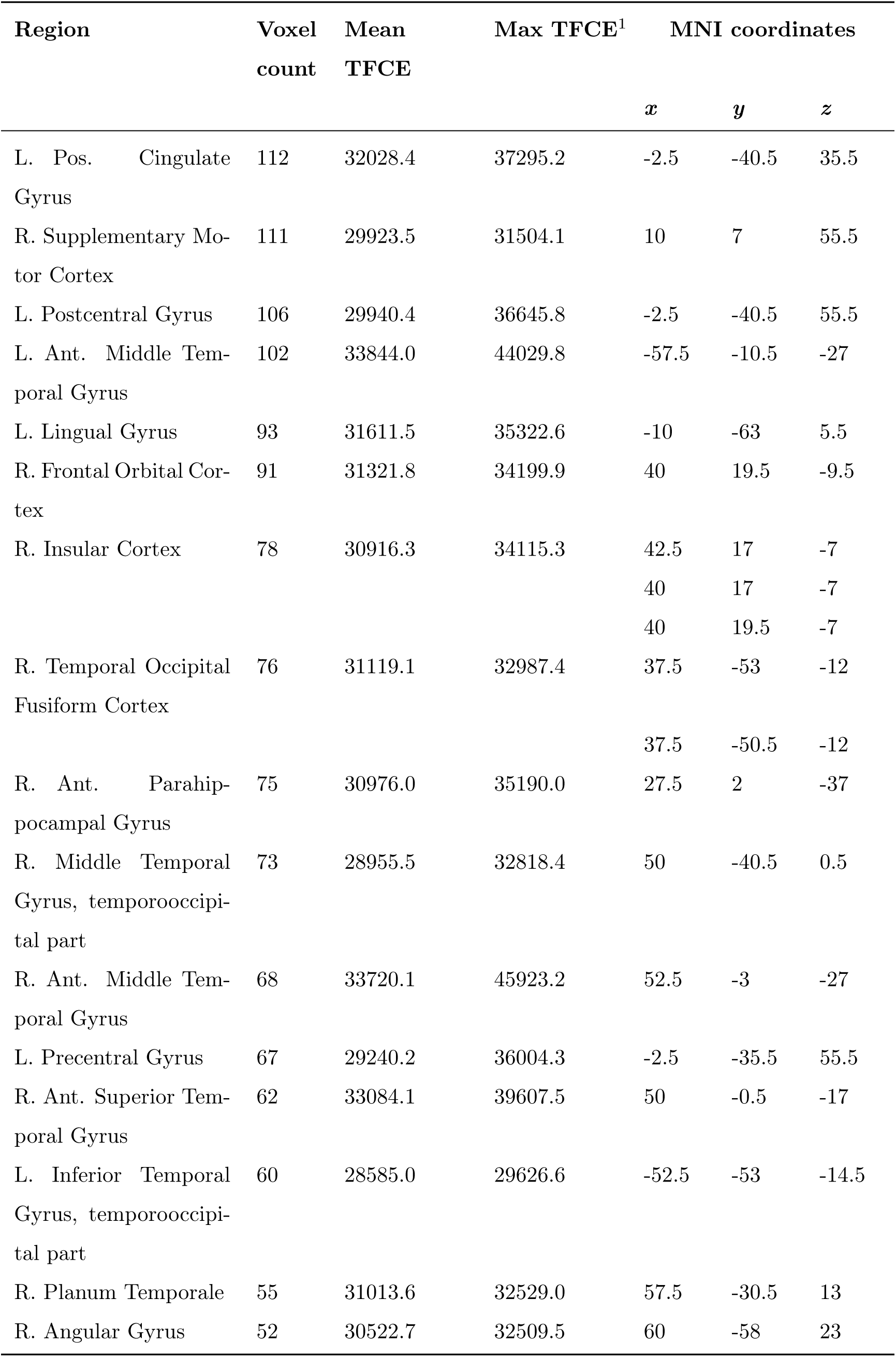
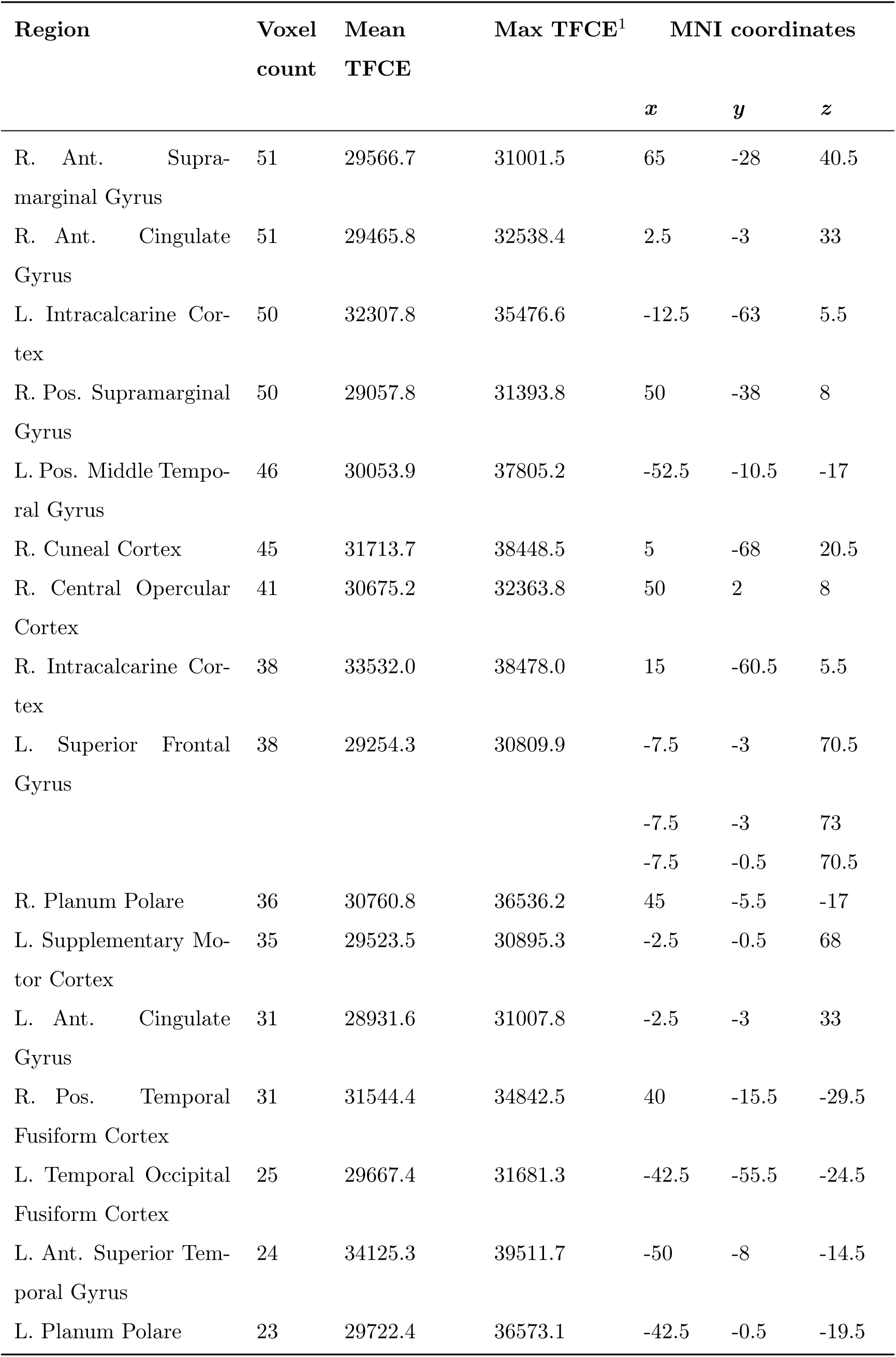
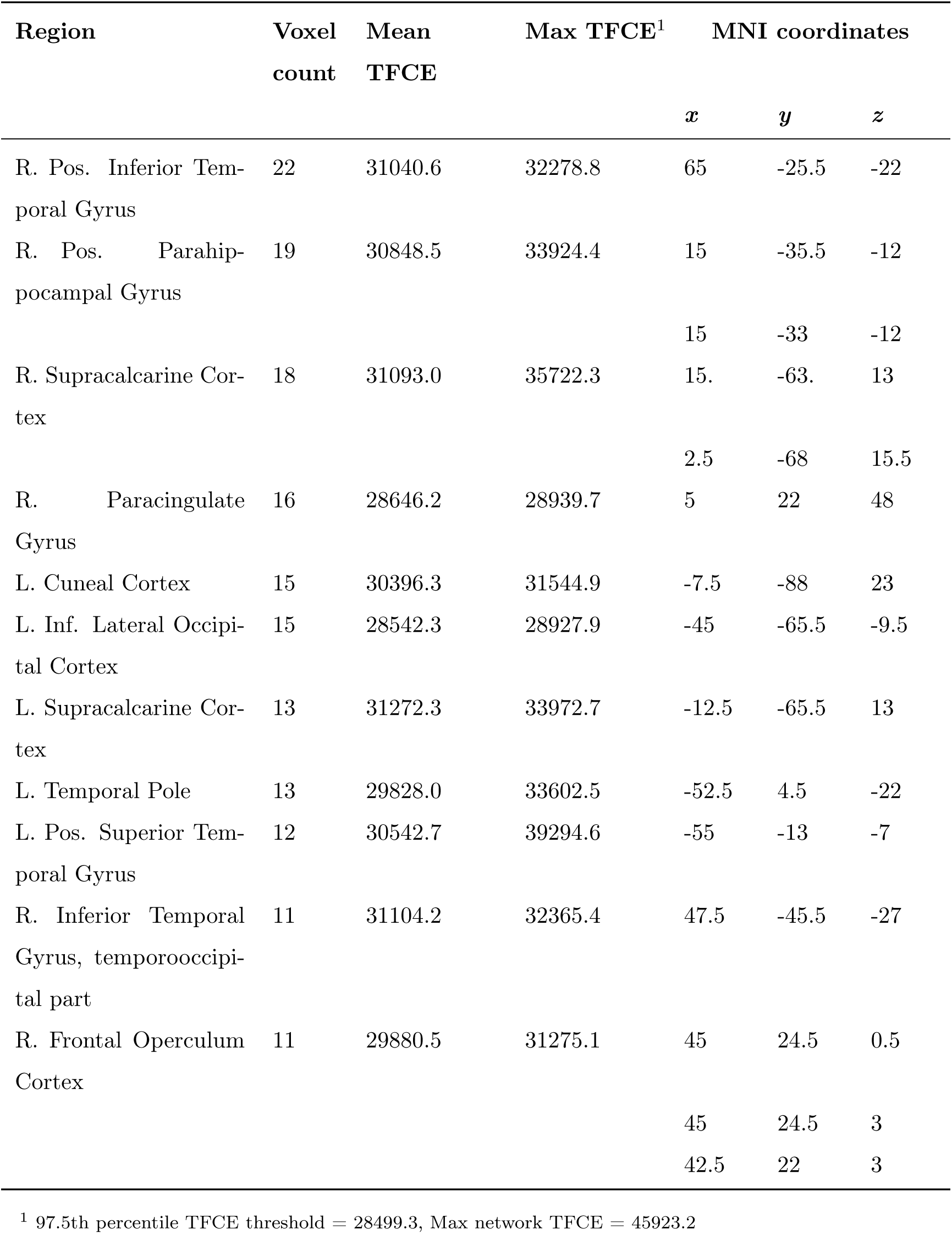
Peak voxel coordinates of regions with at least 10 voxels in the general personal semantic network (Equation 4, Figure 4A). The FSL-Harvard-Oxford cortical-subcortical atlas was used to get coordinates in MNI space. When multiple sets of coordinates are shown for a region, they correspond to multiple peak voxels.

### S2 List of regions in the vivid personal semantic network

**Table S2.**
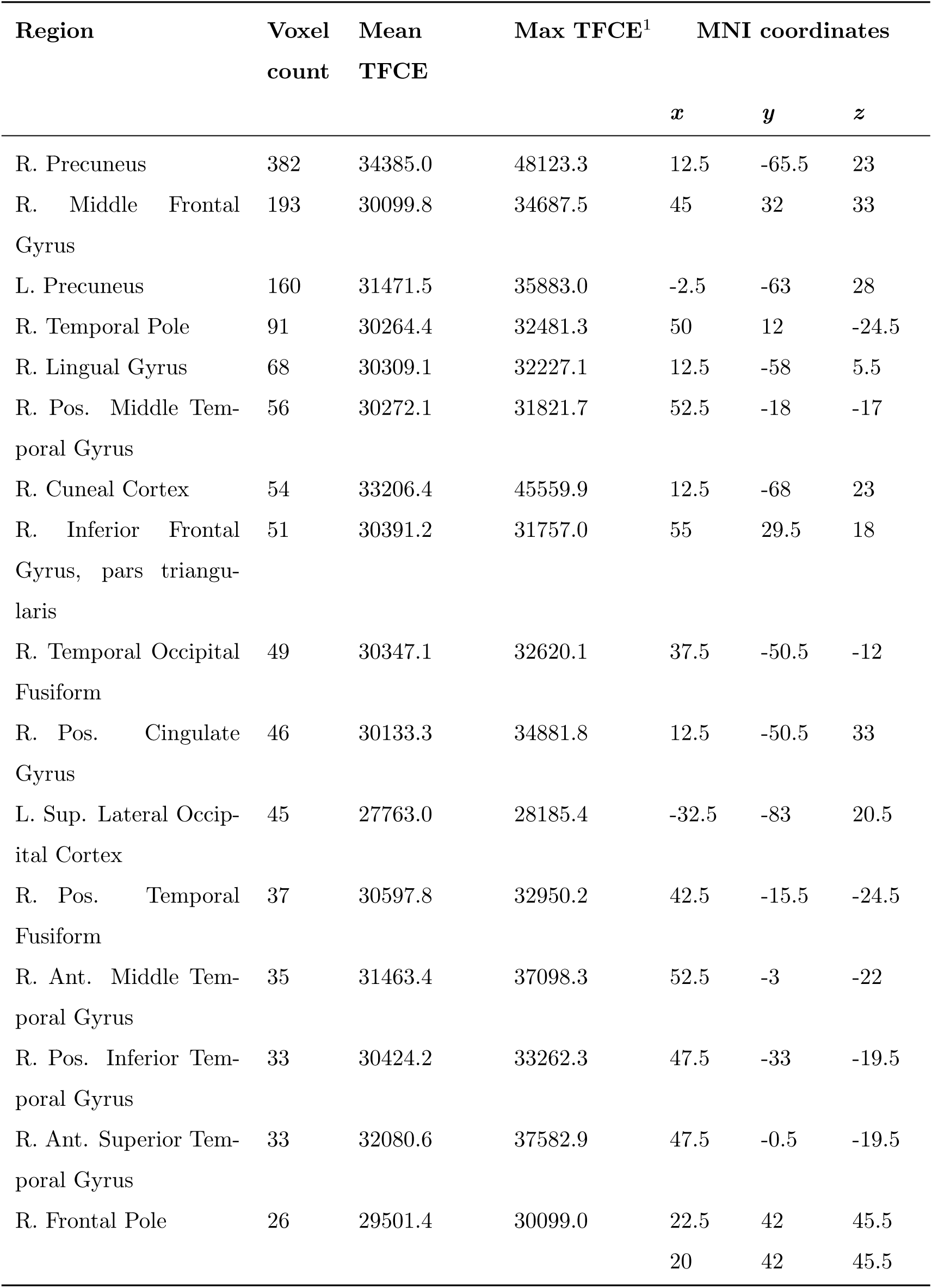
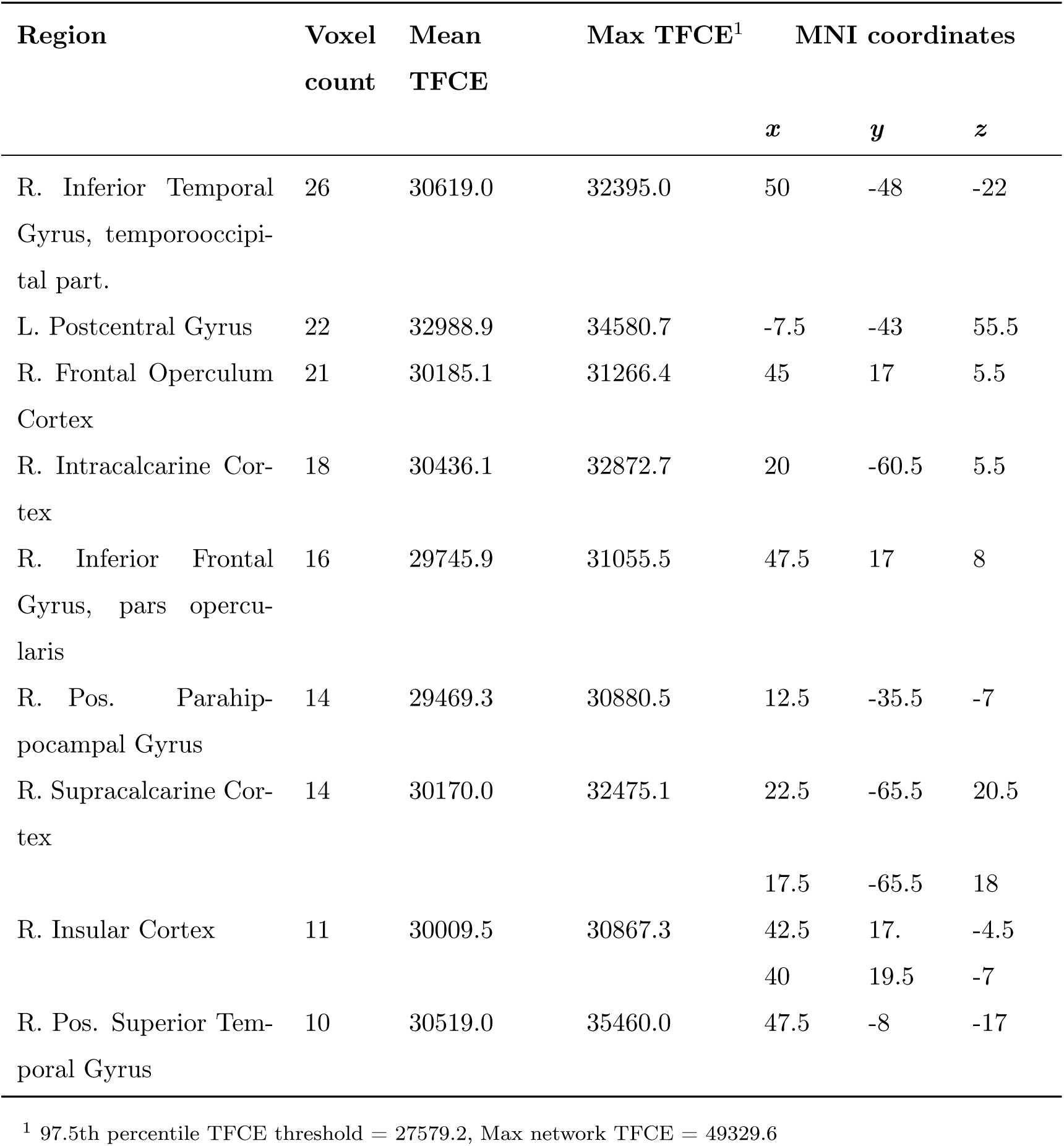
Peak voxel coordinates of regions with at least 10 voxels in the vivid personal semantic network (Equation 5, Figure 4B). The FSL-Harvard-Oxford cortical-subcortical atlas was used to get coordinates in MNI space. When multiple sets of coordinates are shown for a region, they correspond to multiple peak voxels.

## S3 Image Representations

All images were resized to 640 × 480. The color histogram and color correlogram (Huang et al., 1997) representations were computed as in Sreekumar et al. (2014) and descriptions of the methods are reproduced below. Additionally, we explored Histogram of Oriented Gradients (HOG; Dalal et al., 2006; Felzenszwalb et al., 2010, GIST Oliva and Torralba, 2001, and Speeded-up Robust Features (SURF; Bay et al., 2008). SURF is a faster version of the Scale Invariant Feature Transform (SIFT; Lowe, 2004). We provide brief high-level descriptions of each representation below and ask that readers refer to the original papers for more details.

### Color Histogram

The color histogram is a simple global image representation and is invariant under image rotation and translation. A color histogram for an image is generated by concatenating N higher order bits for features in the chosen color space. We used the Hue, Saturation and Value (HSV) space (Smith, 1978) since it separates color from intensity information and makes an image representation based on HSV relatively robust to changes in appearance due to differences in lighting conditions. The histogram is generated by counting the number of pixels with the same color and accumulating it in 2^3*N*^ bins. Quantizing the hue component more precisely than the value and saturation components makes the HSV histogram more sensitive to color differences and less sensitive to brightness and depth differences and could help identify similar images under different lighting conditions. We used a 30 × 10 × 3 hue value saturation quantization of the HSV space to generate 900-dimensional color histogram image vectors.

### Color Correlogram

The color histogram has the drawback of being a purely global description of the color content in an image. It does not include any spatial information. Purely local properties when used can be extremely sensitive to appearance changes due to slight changes in angle, zoom, etc. Purely global properties like those used in the color histograms can give false positives in an image retrieval task as it tends to classify images from widely separated scenes as belonging to the same scene if they have similar color content. A color correlogram describes global distributions of local spatial color correlations. We followed the procedure in Sreekumar et al. (2014) to compute the color correlogram as follows. The color correlogram 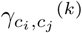 of an image *I*, is a three dimensional table whose entry (*c*_*i*_*, c*_*j*_*, k*) is the probability of finding a pixel of color *c*_*j*_ at a distance *k ∈* {1, 2, 3*,…, d*} from a pixel of color *c*_*i*_ in the image. For pixels *p*1 = (*x*1*, y*1) and *p*2 = (*x*2*, y*2), we use the *L***_*∞*_** norm to measure the distance between them, such that *|p*1 *-p*2*|* = *max*(*|x*1 *-x*2*|, |y*1 *- y*2*|*). Relative to the histogram, the correlogram is robust to changes in appearance caused by occlusions, zoom, and viewing angles. The size of the correlogram is *O*(*m*^2^*d*) where m is the total number of colors and *i, j ∈* {1, 2, 3*,…, m*}. This imposes substantial storage requirements for large values of *d*. So we chose to work with a compressed version of the color correlogram where we sum the conditional probabilities of color pairs over a restricted set of distances. For constructing the color correlograms, the HSV color space is quantized into 12 × 3 × 3 bins. We let *k ∈* {1, 3, 5, 7} and use a restricted version of the color correlogram as in Equation 6.

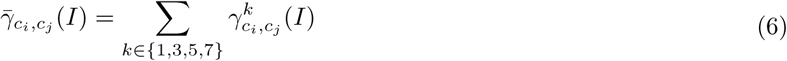

This procedure resulted in 11664-dimensional feature vectors. A singular value decomposition (SVD) is carried on the 120 × 11664 image by feature matrix to generate 120-dimensional image vectors.

### Histogram of Oriented Gradients (HOG)

The histogram of oriented gradients (HOG) is used widely in object detection applications (Dalal et al., 2006). We used the UOCTTI variant (Felzenszwalb et al., 2010) as implemented in VLFeat version 0.9.20 (www.vlfeat.org;Vedaldi and Fulkerson, 2010). HOG computes a histogram of oriented gradients over square cells, typically 8 pixels perside. We also used the typical value of orientation bin size of 9. Since images were 640 × 480 pixels, there were 80 × 60 HOG cells. Dalal et al. (2006) originally proposed normalization and truncation of HOG features via 4 normalization factors to obtain 36 HOG features. Felzenszwalb et al. (2010) proposed alternative steps (using fewer features to speed up learning and detection) involving a principle components analysis of a collection of HOG features to derive 13 contrast-insensitive HOG features. However, their analyses also indicated that detection performance for some object classes improved when using some additional contrast-sensitive features. The end result is a 31-dimensional feature vector (see Felzenszwalb et al., 2010 for further details). Thus, we obtain 80 × 60 × 31 = 148800-dimensional HOG vectors. A singular value decomposition (SVD) is carried on the 120 × 148800 image by feature matrix to generate 120-dimensional image vectors.

### GIST

Oliva and Torralba (2001) proposed a model of real-world scene recognition, based on a low dimensional scene representation that they called “Spatial Envelope”. Unlike HOG, this model was not designed to detect individual objects, rather it was aimed at representing dominant spatial characteristics of a scene using a set of perceptual dimensions that were estimated using spectral and coarsely localized information. This model successfully models a holistic representation of the scene and generates a multidimensional space in which semantic categories of scenes (e.g. highways) cluster together. MATLAB code provided by the original authors was used to construct GIST representations for our analyses (http://people.csail.mit.edu/torralba/code/spatialenvelope/).

### Speeded-up Robust Features (SURF)

Speeded-up Robust Features (SURF), as the name suggests, is a fast algorithm partly inspired by the Scale Invariant Feature Transform (SIFT), that detects interest points in a view-invariant manner. We first used the bagofFeature() function in the MATLAB computer vision system toolbox to extract SURF features for all 120 images for a given participant. As an example, 1843200 features were extracted from the image set for one participant. K-means clustering is performed to create a 500-word visual vocabulary. Each image is then represented as a histogram over these 500 clusters using the MATLAB computer vision system toolbox *encode()* function.

## S4 Comparing image representations: Common neighbor ratio

As in Sreekumar et al. (2014), to pick the “best”image representation for subsequent analyses, we required that our representation of choice and the associated distance measure accurately identify images from the same context as being similar to each other. Though people can and do go back to the same context at a later time, in general images that are close in time will be from the same spatial context and hence should be identified as being similar. With this in mind, we defined the common neighbor ratio (CNR). Given a positive integer *k*, for each image *I*, we find its *k* nearest neighbors both in the spatial domain and in the time domain. Suppose *D*_*I*_ = {*I*_*d*1_*, I*_*d*2_*, I*_*d*3_*,…, I*_*dk*_} are image *I*’s *k* nearest neighbors in space and *T*_*I*_ = {*I*_*t*1_*, I*_*t*2_*, I*_*t*3_*,…, I*_*tk*_} are image *I*’s *k* nearest neighbors in time, then

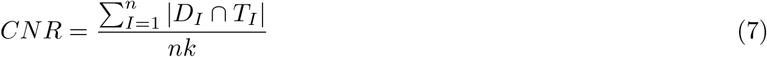

where *n* is the total number of images. If *k* equals *n -* 1 (i.e., all the other images in the set), then the ratio is1. The method that has a higher common neighbor ratio is the better one for our purpose, which is to successfully identify images that came from the same context as similar. Figure S1 shows common neighbor ratios averaged over participants for each image representation. For all reasonable values of *k* nearest neighbors (given that care was exercised while selecting stimuli to avoid images that came from the same episode too often, it is unlikely that there are many neighbors from the same temporal context in any given stimulus set of 120 images and so we explored values of *k* up to 15), we found that the color correlogram achieves better congruence between spatial (in image space) and temporal proximity and hence chose the color correlogram as our preferred image representation (Figure S1).

**Figure S1.**
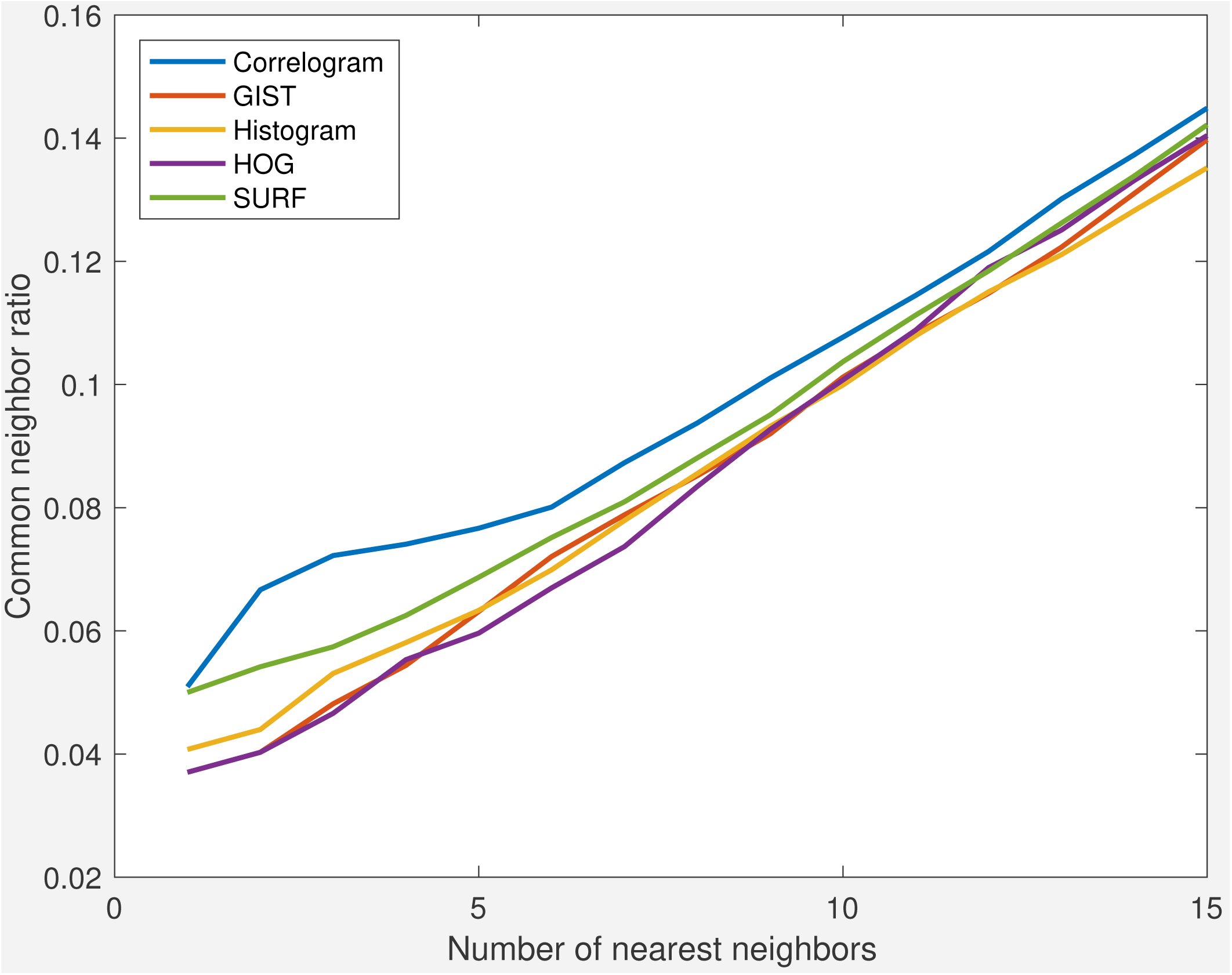
Common neighbor ratio comparison of image representations. The color correlogram representation achieves the best congruence between spatial (in image space) and temporal proximity of nearest neighbors (k).

## S5 Individual Partial Residual Plots

**Figure S2.**
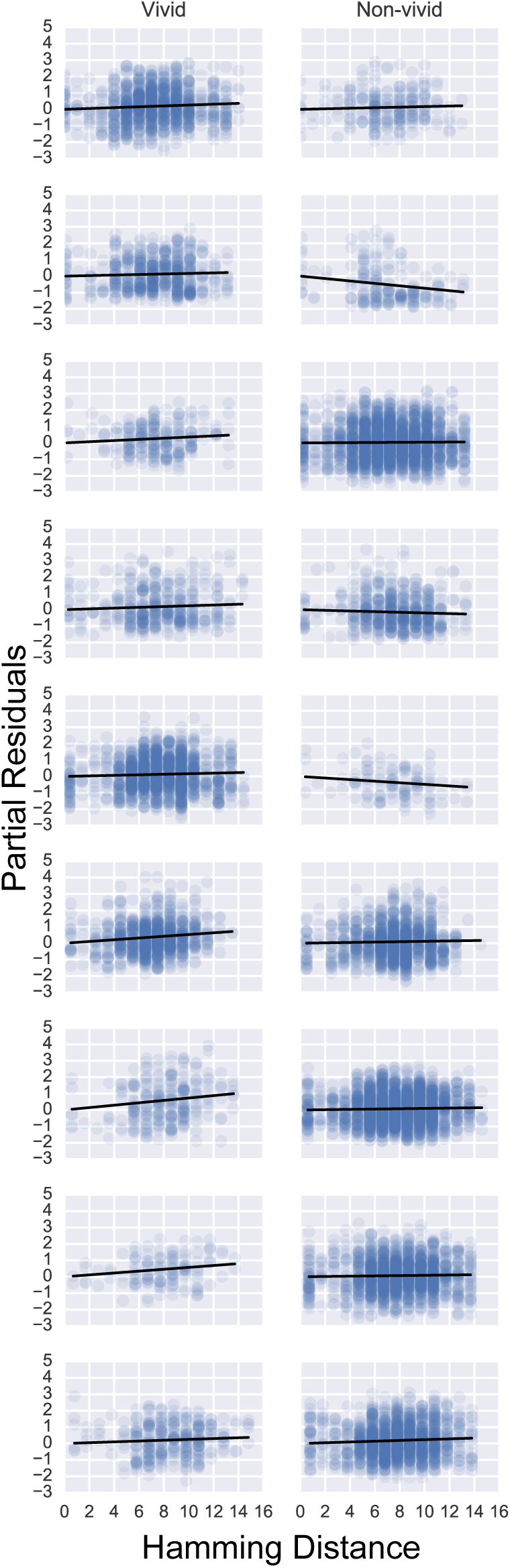
Individual participant partial residual plots of the Neural distance∼ Hamming distance relationship for vivid (left panel) and non-vivid (right panel) pairs

